# Culturomics of *Andropogon gerardii* rhizobiome revealed nitrogen transforming capabilities of stress-tolerant *Pseudomonas* under drought conditions

**DOI:** 10.1101/2022.07.18.500515

**Authors:** Soumyadev Sarkar, Abigail Kamke, Kaitlyn Ward, Eli Hartung, Qinghong Ran, Brandi Feehan, Matthew Galliart, Ari Jumpponen, Loretta Johnson, Sonny T.M. Lee

## Abstract

**Background:** Climate change will result in more frequent droughts that impact soil-inhabiting microbiomes in the agriculturally vital North American perennial grasslands. In this study, we used the combination of culturomics and high-resolution genomic sequencing of microbial consortia isolated from the rhizosphere of a tallgrass prairie foundation grass, *Andropogon gerardii*. We cultivated the plant host-associated microbes under artificial drought-induced conditions and identified the microbe(s) that might play a significant role in the rhizobiome of *Andropogon gerardii* under drought conditions.

**Results:** Phylogenetic analysis of the non-redundant metagenome-assembled genomes (MAGs) identified the bacterial population of interest – MAG-*Pseudomonas*. Further metabolic pathway and pangenome analyses detected genes and pathways related to nitrogen transformation and stress responses in MAG-*Pseudomonas*.

**Conclusions:** Our data indicate that the metagenome-assembled MAG-*Pseudomonas* has the functional potential to contribute to the plant host’s growth during stressful conditions. This study provided insights into optimizing plant productivity under drought conditions.

## Background

Global climate change is a serious concern, resulting in soil degradation, soil erosion, and impacts on soil health [1]. Climate change has severe impacts worldwide including in the USA, resulting in more frequent and prolonged droughts [2], and is gradually degrading the plant diversity and ecosystem functions [3]. The rhizobiome, microbial communities that are intimately associated with the plant rhizosphere [4,5], is one of the key factors in maintaining ecosystem function, soil quality and plant health [6,7]. The plant rhizosphere is the primary site for plant-microbe and microbe-microbe interactions, governed primarily by root exudates [8]. Microbes in the rhizobiome can facilitate plant host nutrient and water uptake, element cycling (carbon, nitrogen, phosphorus) and other processes that are beneficial to plants [9–11].

Rhizobiomes are also instrumental in enhancing plant hosts’ resistance and resilience against abiotic stresses such as drought, salinity, and heavy metal exposure [12]. Therefore, with the more frequent and more extreme droughts events predicted in the global climate change scenarios in the future, it is even more urgent to provide insights into the mechanisms of how the rhizobiome may promote plant host resilience and response to stress. Although there are various studies that have dissected how climate change impacts the rhizobiome [13–16], more concerted efforts are needed to provide insights into the mechanisms of how the rhizobiome can enhance the plant host resilience during drought-induced experimental stress.

Previous studies have reported a clear contribution from plant-associated microbial members to plant growth and resilience during drought conditions [17–19]. Plant growth-promoting bacteria (PGPB) reportedly enhance plant growth during drought [20,21], an observation attributed to the microbial nitrogen cycling and transformation in soil [22]. Therefore, candidate microbes capable of nitrogen transformation and increase nitrogen availability in the rhizosphere have been the key targets in a growing number of experimental and observational studies that focus on the assembly of plant health promoting Synthetic Communities (SynCom) [23,24]. SynComs have been successfully deployed to alter the plant phenotype, to enhance plant disease resistance and productivity [25,26]. However, it is challenging and tedious to select optimal members of SynComs because of the lack of knowledge of the microorganisms that could impart favorable functions under stressful conditions [27]. Therefore, in identifying candidate microbes for SynComs, it may be more expedient to identify specific microbial functions and mechanisms rather than to depend solely on taxonomy.

Our long-term, ongoing research on the microbiome of dominant Great Plains prairie grass *Andropogon gerardii* (Big Bluestem) provided an excellent opportunity to acquire deeper insights into the microbial functional potential under abiotic stress [28–30]. There are three *A. gerardii* ecotypes (dry, mesic, and wet) that originated in Hays, Kansas (rainfall ∼500 mm/year), Manhattan, Kansas (rainfall ∼870 mm/year) and Carbondale, Illinois (rainfall ∼1,200 mm/year), respectively [28,29]. In this study, we attempted to elucidate the rhizosphere microbial functional potential from *A. gerardii* growing in Colby, Kansas, where the low precipitation defines a margin of the environment suitable for *A. gerardii* survival and growth. We aimed to identify the *A. gerardii* rhizobiome associated microbial population(s) that are drought resistant or resilient, and to acquire insights into the microbial functions by: (1) isolating and cultivating microbes that existed in the ecotypes using media that promote drought-induced stress [28]; (2) obtaining genomic insights into drought resilient bacterial populations that can contribute to the nitrogen transformation. In this study, we combined culturomics and high-resolution genomic sequencing to identify microbial populations and their functional potential to enhance *A. gerardii* resistance and resilience during drought stress.

## Results and discussion

### MAGs analysis, phylogenetic analysis, and identification of MAG-Pseudomonas

We cultured the *A. gerardii* rhizosphere microbial populations using the following samples and media - dry ecotype in R2A, dry ecotype in R2A with PEG, wet ecotype in R2A, and wet ecotype in R2A with PEG. We expected that the PEG-amended media would yield bacterial populations enriched with drought resistant gene functions. We recovered a total of 125 MAGs and generated a total of 63 non-redundant MAGs from the four conditions (Figure 1A, Supplementary Table S1). We recovered an average of 173,480 ± 22,383 contigs, with N50 of 33,485 ± 2,526. The resolved MAGs had completion values of 89.4% ± 2.0%, and ∼92.8% were annotated to the genus level (Supplementary Table S1). Among the 63 non-redundant MAGs, the dominant phyla were Proteobacteria (n=20), Firmicutes (n=36) and Actinobacteria (n=7).

**Figure 1:**
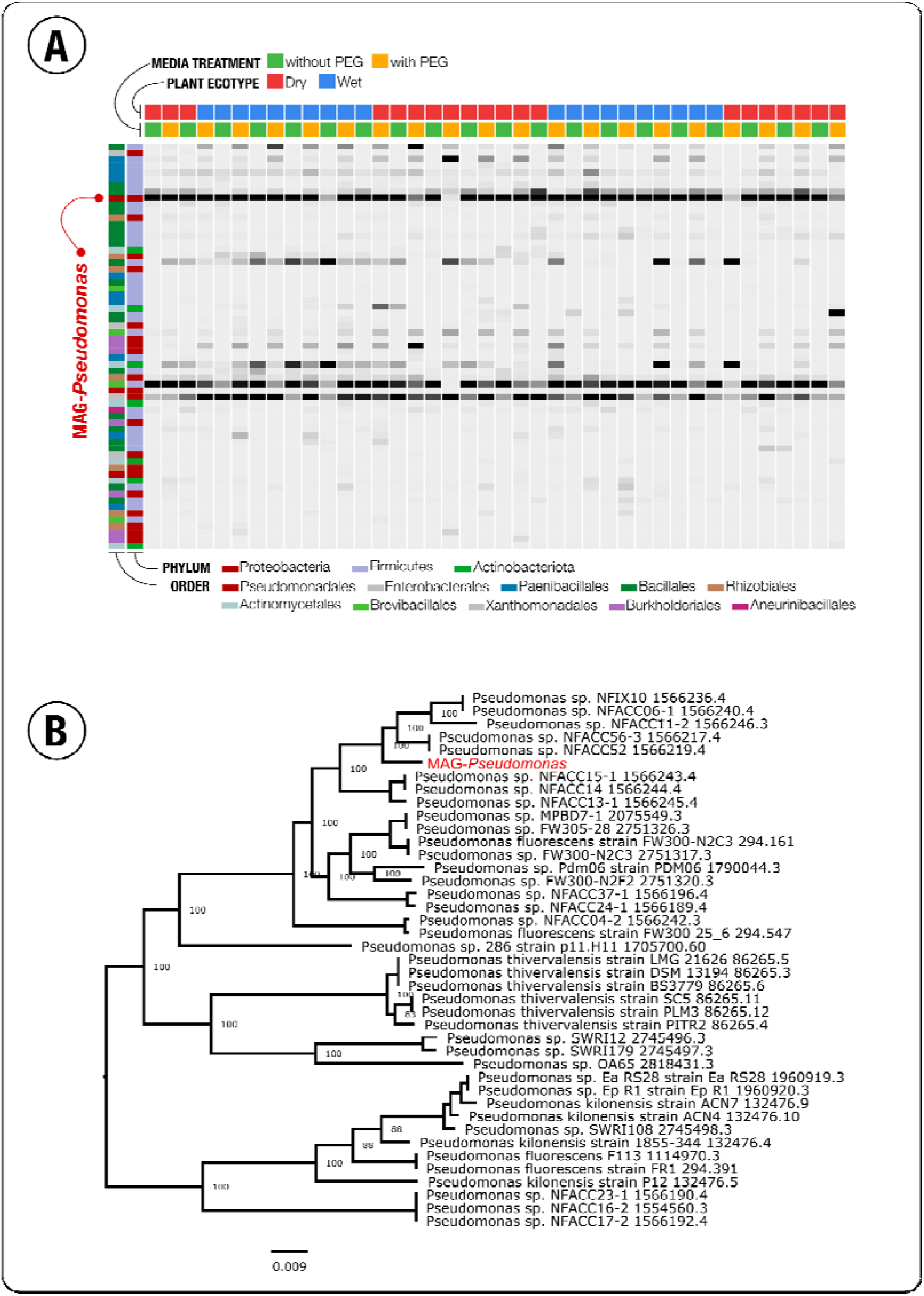
(A) Detection of non-redundant metagenome-assemble genomes (MAGs) in the rhizosphere of dry and wet *Andropogon gerardii* ecotypes when cultivated in normal precipitation (without PEG) and under drought-induced conditions (with PEG). MAG-*Pseudomonas* was highly detected in all growing media conditions of both dry and wet ecotypic rhizosphere samples. (B) Phylogenetic analysis of MAG-*Pseudomonas* with closely related 40 genomes. *Pseudomonas* sp. NFACC52 is the most closely related genomes to MAG-*Pseudomonas*.

Among the 63 non-redundant MAGs that we resolved, one of the clusters (consisting of four MAGs having > 95% ANI identity) was assigned to the genus *Pseudomonas*. We obtained 40 closest related *Pseudomonas* genomes using the Similar Genome Finder service, and confirmed that the MAG had the greatest similarity to *Pseudomonas sp*. NFACC52 (Figure 1B, Supplementary Table S1). We observed that the representative MAG (MAG_001; hereafter referred to as MAG-*Pseudomonas*) for this cluster was highly detected in all the culture conditions, and with their ubiquitous presence in the soil irrespective of the ecotype and drought stress, we hypothesized that MAG-*Pseudomonas* might be an important contributor in the rhizobiome associated with *A. gerardii. Pseudomonas spp*. are common in the rhizosphere and reported to have important functions in modulating host performance [54–56]. *Pseudomonas thivervalensis* were isolated from the roots of *Brassica napus* and *Arabidopsis thaliana [57]*, and is a functionally significant member of soil microbial communities [55]. *Pseudomonas* have also been implicated to be a plant growth-promoting rhizobacteria (PGPR) and have been associated with plant growth, control of pathogenicity [55] and aid in plant resilience under drought-stressed conditions [54,56].

### Stress response genes identified in MAG-Pseudomonas enhanced drought tolerance

MAG-*Pseudomonas* has a total length of 6,777,975 bp, with 99 contigs and an N50 of 146,692 bp. The GC content of MAG-*Pseudomonas* is 61.1%. When annotated with the COG database, we noticed that it yielded 5,953 gene calls, and 4,924 were assigned at least one COG categorical function (Table 1).

**Table 1:**
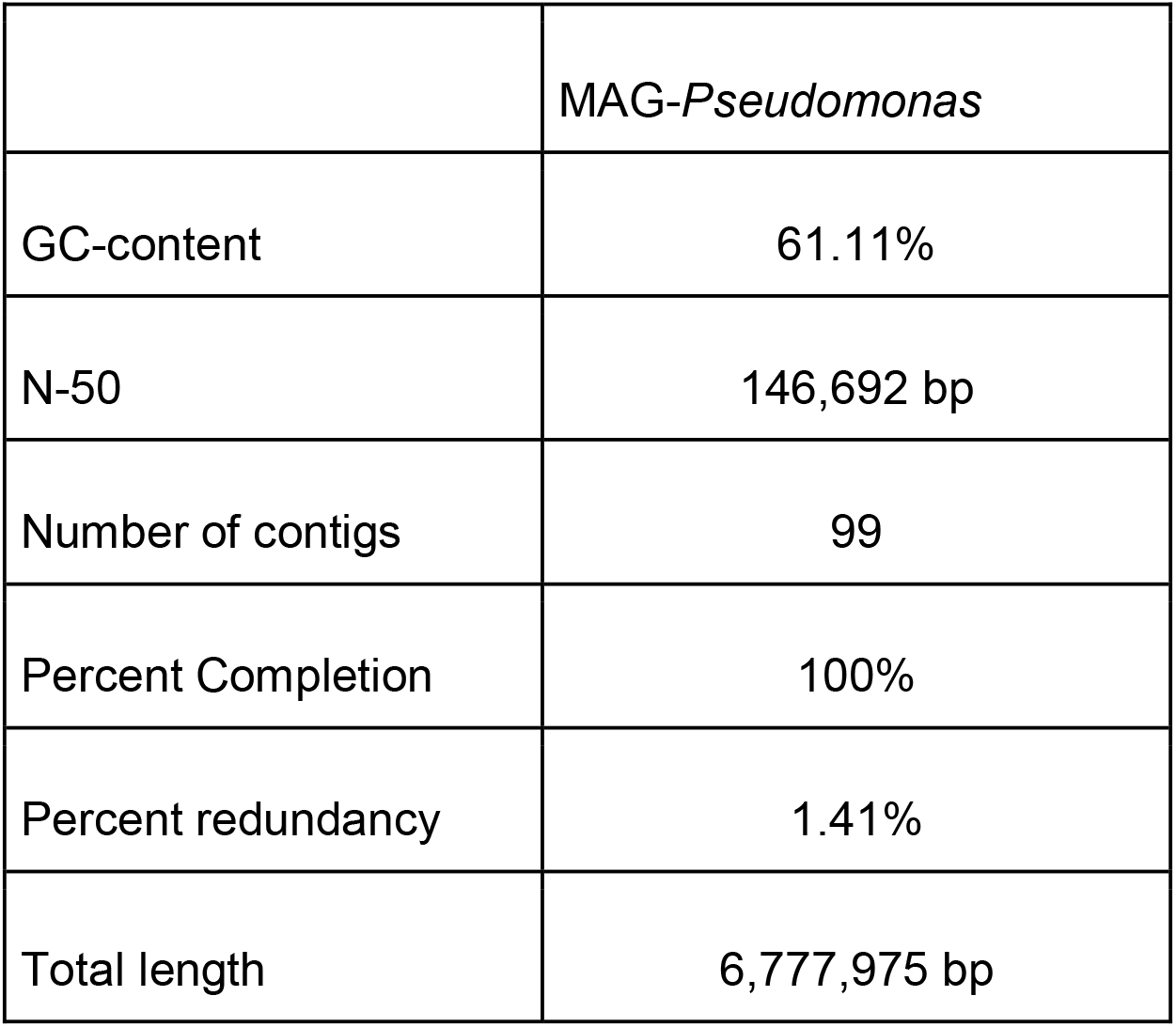
Assembly statistics of MAG-*Pseudomonas*: GC-content, N-50, number of contigs, percent completion, percent redundancy, and total length.

Previous studies [54,56] based on 16S rRNA gene sequences have identified *Pseudomonas* in aiding the plant host to become more resilient under drought-stressed conditions. This led us to ask what might be some potential functions of *Pseudomonas* that 1) enabled the survival of *Pseudomonas* under stressful conditions; 2) provided cues to how the *Pseudomonas* might assist in the stress tolerance of the associated plant host. We observed 14 COG functions that were assigned to stress responses, with 6 gene functions that were in the universal stress protein (UspA) family, that might enhance the survivability of *Pseudomonas* during stressful conditions. There were 3 COG functions that were assigned to Ser/Thr protein kinase RdoA which was involved in the Cpx stress response. The other gene functions that were identified related to stress responses were putative negative regulator of RcsB (n=1), acid stress-induced BolA-like protein IbaG/YrbA (n=1), stress-induced morphogen (n=1), ribosomal protein L25 (n=1), various environmental stresses-induced protein Ves (n=1) and predicted membrane GTPase involved in stress response (n=1) (Figure 2; Supplementary Table S2). Universal stress protein (USP) superfamily has an important role in the survivability of bacteria under a wide range of stress conditions [58]. In our study, the amendment of PEG in R2A for cultivation not only mimicked the drought conditions, but also generated moderate levels of osmotic shock for the cells [59], the conditions that often accompany drought conditions [60]. Furthermore, we identified Ves proteins during drought/ osmotic stress in MAG-*Pseudomonas*. Our results suggest that MAG-*Pseudomonas* had the potential to be more resilient and tolerant against drought stress, demonstrating moderate tolerance to dehydration and water limiting conditions [61–64]. Under stress, most microbes will restructure their metabolism and specifically activate various stress pathways [17]. *Pseudomonas* can utilize a range of mechanisms such as alginate [65] and trehalose production [66] to be more resilient during drought conditions. Besides the 14 stress related gene functions, we also identified DNA-binding transcriptional regulator YbjK, (Figure 2; Supplementary Table S2) which is involved in the regulation of stress response [67]. Drought conditions often result in the generation of oxidative stresses in both plants and their associated microbes [68,69]. There is established documentation about the beneficial role of *Pseudomonas* to alleviate the oxidative stress [70,71]. Our results illustrated that our MAG-*Pseudomonas* had several regulatory responses that might enhance its drought stress resilience and fitness.

**Figure 2:**
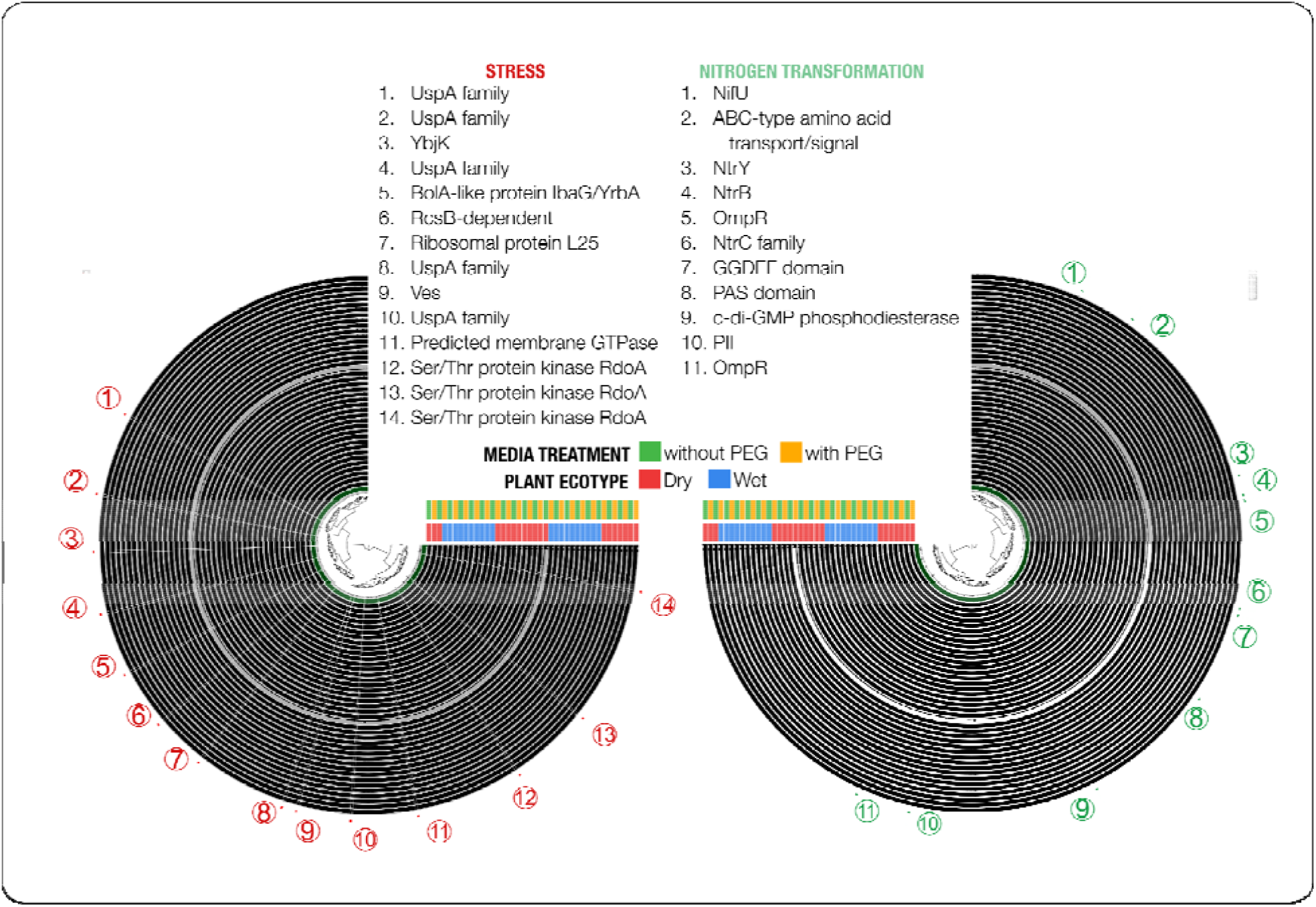
Stress response and nitrogen transformation genes identified in MAG-*Pseudomonas*. Each layer represents a sample (ecotype x media used) and each split represents a contig. There were 14 genes annotated to stress responses and 11 genes were annotated to nitrogen transformation capability.

### Nitrogen transformation potential of MAG-Pseudomonas could enhance A. gerardii growth

Initial genomic analysis showed that our resolved MAG-*Pseudomonas* harbored several stress-response related gene functions. Besides understanding microbial mechanisms of MAG-*Pseudomonas* resilience during drought-induced stress, we were also interested in gaining a deeper understanding of how the plant host could benefit from the *A. gerardii* and MAG-*Pseudomonas* interactions. We detected several gene functions that demonstrated the nitrogen transformation potential in our resolved MAG-*Pseudomonas*, which could contribute to the growth enhancement of the associated plant host, *A. gerardii*.

We detected a number of gene functions that are related to nitrogen transformation - nitrogen fixation protein NifU and related proteins; signal transduction histidine kinase NtrY involved in nitrogen fixation and metabolism regulation (NtrY); signal transduction histidine kinase NtrB, nitrogen specific (NtrB), two-component system; NtrC family (nitrogen regulation response regulator GlnG, nitrogen PTS system EIIA component, and nitrogen regulatory protein PII, GlnK) (Figure 2, Supplementary Table S2). These genes have been previously reported in *Pseudomonas spp [72–74]*. All the nitrogen transformation gene functions that were detected in our MAG-*Pseudomonas* can be essential in helping to fulfill the plant host’s need for nitrogen, especially in N-depleted soils [75–78]. Bacterial Nif genes transform the atmospheric nitrogen to the form that can be utilized by the plants [79,80], whereas NtrY encodes for the sensory kinase of the two-component regulatory system of NtrY/NtrX associated with nitrogen metabolism [81]. NtrB also plays a role in nitrogen metabolism and can regulate the nitrogen dynamics under nitrogen-deprived and enriched environments [82]. NtrC is another nitrogen metabolism regulator that contributes to nitrogen assimilation [73]. Similarly, nitrogen regulatory protein PII (GlnK) and nitrogen PTS system EIIA components are also involved in regulating nitrogen metabolism [83]. Besides the Nif genes, we further detected gene functions in the MAG-*Pseudomonas* that corresponded to nitrogen assimilation and nitrogen dissimilation (nitrification and denitrification) (Figure 2, Supplementary Table S2), which contributes to the nitrogen cycle [84]. Assimilatory nitrate reductase catalytic subunit was identified in this study that catalyzes the process from nitrate to nitrite [85]. NAD(P)H-nitrite reductase, a large subunit (NirB) was detected that can catalyze nitrite reduction, and forms ammonia [86]. We also detected nitrite reductase (NADH) large and small subunits that can carry out similar processes and contribute to the nitrogen cycle [86].

We used the comparative pathway tool in PATRIC, and identified 138 potential pathways of MAG-*Pseudomonas* based on genomic information from 3 *Pseudomonas* genomes - *Pseudomonas chlororaphis subsp. aurantiaca strain* ARS 38 isolated from the cotton rhizosphere, *Pseudomonas sp*. DR208 and *Pseudomonas sp*. DR48 isolated from the soybean rhizosphere. The identified pathway classes included carbohydrate metabolism, lipid metabolism, metabolism of cofactors and vitamins, energy metabolism, nucleotide metabolism, biosynthesis of secondary metabolites, amino acid metabolism, xenobiotics biodegradation and metabolism, metabolism of other amino acids, glycan biosynthesis and metabolism, translation, signal transduction, and immune system (Supplementary Table S3). We further analyzed the differential occurrence of the genes in MAG-*Pseudomonas* and the 3 *Pseudomonas* genomes, and observed that there was a higher occurrence of nitrate reductase, nitrate reductase (NO-forming), and ubiquinol-cytochrome-c reductase in the MAG-*Pseudomonas* than in the other 3 *Pseudomonas* genomes (Figure 3A). Given the importance of nitrogen-mediated microbial processes on plant growth [87], we analyzed the annotated MAG-*Pseudomonas* nitrogen transformation processes and showed that there were 79 annotated genes that were involved in the nitrogen metabolism pathway with 100% coverage. We identified several important genes in MAG-*Pseudomonas* associated with nitrogen transformation processes such as nitrate reductase (n=14), nitrate reductase (NO-forming) (n=6), ubiquinol-cytochrome-c reductase (n=4), NADH:ubiquinone reductase (H (+) - translocating (n=2), cytochrome-c oxidase (n=2), carbamate kinase (n=2), glutamin-(aspargin-)ase (n=2), formate dehydrogenase (n=1), glutamate synthase (NADPH) (n=1), glutamate dehydrogenase (n=1), glutamate dehydrogenase (NADP(+)) (n=1), glutamate synthase (ferredoxin) (n=1), aminomethyltransferase (n=1), asparginase (n=1), glutaminase (n=1), nitrilase (n=1), carbonate dehydratase (n=1), cyanase (n=1), aspartate ammonia-lyase (n=1), glutamate-ammonia ligase (n=1), and aspargine synthase (glutamine-hydrolyzing) (n=1) (Figure 3B). We observed a high number of genes that were related to nitrate reductase, suggesting its importance to the nitrogen metabolism in our MAG-*Pseudomonas*. The nitrate reductase is critical in reducing nitrate to nitrite for several crop plants, as this reaction leads to the production of proteins that are necessary to maintain plant health [88]. Glutamate synthases are actively involved in ammonia assimilation pathways in bacteria [89], while glutamate dehydrogenase has a prominent role in nitrogen assimilation and is capable of maintaining the balance of carbon and nitrogen [90]. Putting it all together, our resolved MAG-*Pseudomonas* with its potential in microbial-driven nitrogen transformation processes could play a critical role in the regulation of primary productivity of its plant host, *A. gerardii*, even during times of drought-induced stress.

**Figure 3:**
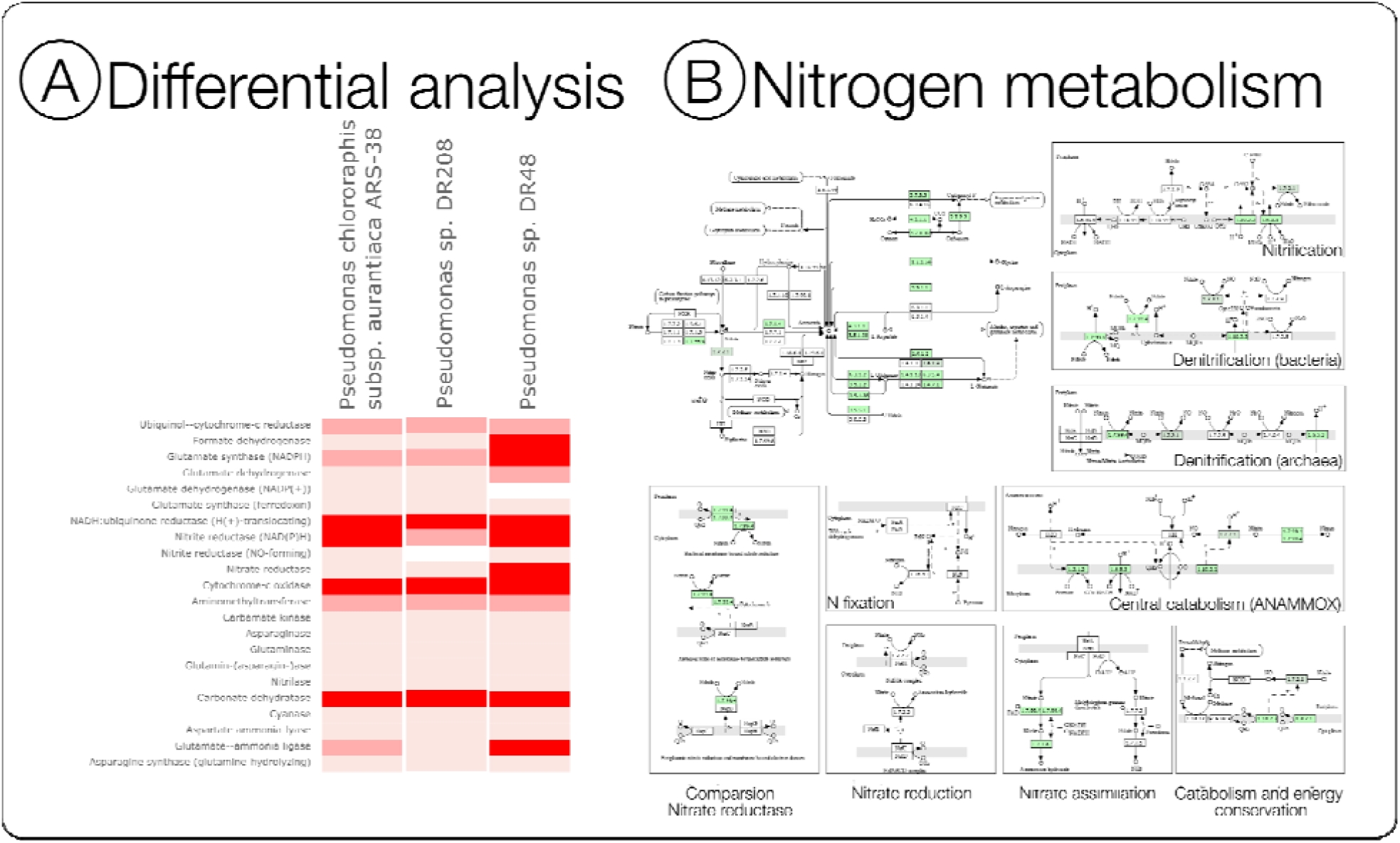
(A) Differential occurrence of the genes in MAG-*Pseudomonas* with *Pseudomonas chlororaphis* subsp. *aurantiaca strain* ARS 38, *Pseudomonas sp*. DR208 and *Pseudomonas sp*. DR48. The darker the highlight represents higher occurrences in the MAG-*Pseudomonas*. MAG-*Pseudomonas* showed high differential occurrences in nitrate reductase, nitrate reductase (NO-forming), and ubiquinol-cytochrome-c reductase as compared to the other 3 genomes. (B) Nitrogen metabolism pathways in MAG-*Pseudomonas* were detected based on a comparative pathway tool in PATRIC. MAG-*Pseudomonas* had 79 annotated genes involved in the nitrogen metabolism pathway with 100% coverage.

### MAG-Pseudomonas is essential to understand the resilience of the host plant under abiotic stress

Our genomic analysis revealed several stress response and nitrogen transformation functional potential, but are there any niche specificity for MAG-*Pseudomonas*? We used a pangeomic analysis to compare the shared and unique gene functional potential of MAG-*Pseudomonas* and 6 closely-related *Pseudomonas* genomes. Our analysis yielded 39,798 genes across the 7 genomes, with a total of 12,473 gene clusters. We used hierarchical clustering to group the gene clusters, showing similar distribution patterns across the 7 genomes (Figure 4, Supplementary Table S4). Our pangenomic analyses identified a collection of 15,605 core gene clusters that occurred in all *Pseudomonas* genomes, and 752 gene clusters that only occurred in the MAG-*Pseudomonas* genome. The proportion of genes with functional annotation varies between the core and accessory clusters of MAG-*Pseudomonas*. We noticed that there were 94.4% of core gene clusters annotated with gene functions, using NCBI’s Clusters of Orthologous Groups (COGs) database, while only 63.6% of the gene clusters in the accessory clusters of MAG-*Pseudomonas* were annotated (Figure 4, Supplementary Table S4).

**Figure 4:**
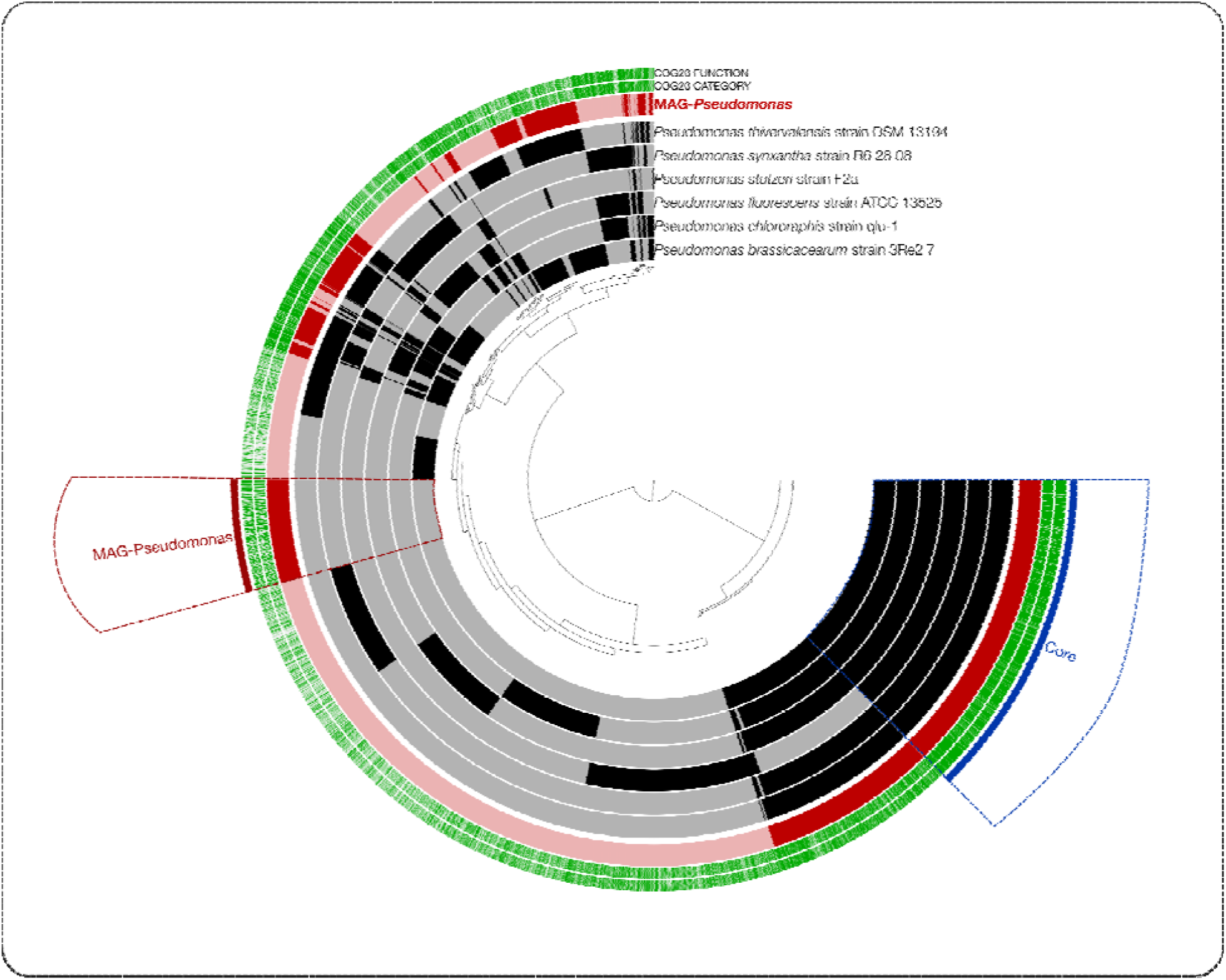
Pangenomic analysis of MAG-*Pseudomonas* with 6 closely-related *Pseudomonas* genomes. The closely-related genomes include *Pseudomonas thivervalensis* strain DSM 13194, *Pseudomonas synxantha* strain R6 28 08, *Pseudomonas stutzeri* strain F2a, *Pseudomonas fluorescens* strain ATCC 13525, *Pseudomonas chlororaphis* strain qlu-1, and P*seudomonas brassicacearum* strain 3Re27. The pangenomic analyses show the core gene clusters and MAG-*Pseudomonas* only accessory gene clusters.

We identified several stress response and nitrogen transformation genes in the core cluster in the pangenomic analysis which again reiterated our hypotheses that MAG-*Pseudomonas* might have microbial mechanisms that enhanced its survivability and would contribute to the plant hosts’ well-being under abiotic stress conditions (Figure 4, Supplementary Table S4). The genes that were identified were predicted membrane GTPase TypA/BipA involved in stress response (TypA), putative negative regulator of RcsB-dependent stress response, UPF0070 family (YfgM), universal stress protein E, nucleotide-binding universal stress protein UspA family (UspA), acid stress protein IbaG/YrbA, BolA-like family (IbaG), Ser/Thr protein kinase RdoA involved in Cpx stress response, MazF antagonist (SrkA), desiccation stress tolerance protein with LEA/WHy domain (LEA), BolA family transcriptional regulator, general stress-responsive regulator, uncharacterized stress-responsive protein, DNA-binding protein YaaA associated with the oxidative stress response (YaaA), universal stress protein A, and Ribosomal protein L25 (general stress protein Ctc) (RplY) (Figure 4, Supplementary Table S4). Desiccation stress tolerance proteins with LEA/WHy domain (LEA) is suggested to confer a broad range of stress response function to bacteria such as *Escherichia coli [91]*, while genes corresponding to a WHy protein homologue have been identified in both archaea and bacteria including *Pseudomonas [92*,*93]* although the specific function in *Pseudomonas* is still incomprehensible. Our findings in MAG-*Pseudomonas* and 6-closely related genomes provided insights into potential gene functions in *Pseudomonas* that could be instrumental in providing resilience against drought induced stress. We also identified a set of universal stress proteins (UspA, UspE), which belonged to bacterial universal stress proteins, and were produced under stressful conditions [94]. We also identified other genes - YaaA (oxidative stress); TypA/BipA (general stress-response regulator, [95]; BolA (family transcriptional regulator, [96]; and Ribosomal protein L25 (general stress protein Ctc) (RplY) [97], that demonstrated the potentiality of *Pseudomonas* to elicit one or more microbial mechanisms to become more resilient when subjected to abiotic stresses. Similar to stress response genes, our findings, which identified numerous nitrogen transformation gene functions (Figure 4, Supplementary Table S4), in the pangenome analysis corroborate with our MAG-*Pseudomonas* genome analysis that *Pseudomonas* might have the capability to contribute to the resilience and well-being of the plant host under environmental stresses.

Besides the core-clusters gene functions, we also observed genes related to chemotaxis in the MAG-*Pseudomonas* accessory gene-clusters. We detected genes corresponding to methyl-accepting chemotaxis protein and chemotaxis protein CheD (Figure 4, Supplementary Table S4). Our resolved MAG-*Pseudomonas* might show chemotaxis towards certain amino acids by using methyl-accepting chemotaxis proteins [98], as these bacterial cells are known to methylate the methyl-accepting chemotaxis proteins when adapting to environmental repellents and attractants [99]. Similarly, CheD chemotaxis proteins might be used by MAG-*Pseudomonas* to attract or evade various environmental stimuli [100–102]. Our MAG-*Pseudomonas* also had a gene corresponding to insecticidal toxin complex protein TccC. These proteins exhibit toxicity to a wide range of insects that could be utilized in designing strategies for crop protection [103]. Interestingly, we also identified the LuxR family transcriptional regulator, quorum-sensing system regulator ExpR in MAG-*Pseudomonas*, suggesting that MAG-*Pseudomonas* might use the LuxR proteins to communicate with neighboring bacteria [104] involving Acyl homoserine lactone (AHL)-dependent Quorum Sensing mechanism [105]. MAG-*Pseudomonas*, therefore, has a plethora of gene functions that might enable the microbe to show phenomena such as chemotaxis and quorum sensing.

Tailoring SynComs is an important approach to provide insights into plant host-microbe interactions. Understanding the mechanisms and functions of host-associated microbial populations is particularly relevant in the construction of these plant-associated SynComs. Our study showed that MAG-*Pseudomonas* not only possessed the resiliency to survive in drought-induced conditions, but were also able to perform essential microbial functions for generating products related to the nitrogen cycle [106], which could be exploited by plant host and other host-associated microbes [107]. A SynCom consisting of six *Pseudomonas* strains isolated from the garlic rhizosphere has been reported to promote plant growth [108]. Apart from the potential to contribute to the plant host’s well-being, our MAG-*Pseudomonas* might also be able to influence and interact with other bacteria, [109], contributing to its role as an important member of the core rhizobiome along with other members such as *Streptomyces, Rhizobium, Burkholderia, Nitrosomonas, Nitrospira, Azospirillum, Bradyrhizobium*, and *Azotobacter* [110]. Overall, our study emphasized that the understanding of the MAG-*Pseudomonas* mechanism and functional potential might contribute to the successful construction of a SynCom that can benefit the plant-host during drought-induced stress [27].

## Conclusion

In this study, we used culturomics and metagenomic strategy to identify bacterial populations in the *A. gerardii* rhizobiome, and identified MAG-*Pseudomonas* as the candidate microbe that had significant functional potential in nitrogen transformation and stress response. In support of other studies, our study verified the abundance of MAG-*Pseudomonas* in the rhizobiome and suggested its potential pivotal role under drought conditions. In a continuing effort to understand the contributions of different microbiota in the plant rhizobiome, it is important to remember that identity and relative abundance alone may not truly reflect the relative functional importance of the bacterial population. Instead, understanding the functional role of the microbe during host-microbe and microbe-microbe interactions might provide more insights. The functional potential of our resolved MAG-*Pseudomonas*, resulting from a combination of conventional culturing and high-throughput analysis, showed the immense potential to inform and refine our efforts to dissect the mechanistic interaction taking place in the rhizobiome.

## Materials and Methods

### Plot design, sampling, and cultivation of rhizosphere communities from soil samples

We collected *Andropogon gerardii* rhizosphere samples from a common garden in Colby at the Kansas State University Agricultural Research Center located in Thomas County (39°23′N, 101°04′W). Further information on the experimental layout, ecotypes, and sampling collections has been described previously [30]. In this comparative study, we selected rhizosphere samples from native dry (Hays, Kansas) and wet (Carbondale, Illinois) ecotypes for microbial cultivation. We separated bulk soil from the soil attached to the rhizosphere by handshaking the roots gently. We dissolved 0.1 g of the rhizosphere samples in 0.9 ml of Phosphate-Buffered Saline (PBS) [pH 7] buffer, serially diluted the samples (10^-1^ - 10^-6^), and spread 100µl solution onto the Petri plates. We designed two culture conditions - 0.315% R2A media (Teknova, USA) [31] and 0.315% R2A media amended with a 36% Polyethylene Glycol 8000 (PEG) (Ψ = -1.54 MPa) to alter the media osmotic potential and to mimic absence and presence of water limitation, respectively [32,33]. A similar range of PEG concentrations has been used to simulate dry environments in other studies [34,35]. To prepare the R2A-PEG media, we 36% (w:v) dissolved PEG powder in autoclaved MilliQ water, allowing the mixed solution (20 mL) to sit on top of a pre-made R2A plate for 24 hours to diffuse throughout the agar. After 24 hours, we removed excess solution and spread 100µl of the diluted soil culture on the surface of the agar. The prepared plates were incubated at 37_ for 24-48 hours until the appearance of the colonies. After the incubation period, we scraped all colonies by flooding the plate with 2 mL of sterile PBS buffer, transferred the liquid that contained microbes and stored at -20□ until genomic DNA extraction. We were interested in capturing the microbial consortia that grew together in different conditions, so instead of picking individual colonies, we scraped all colonies from the individual plates to sequence the full genome(s) [36,37]. Rhizosphere microbial communities were cultivated from dry (R2A; n=10 and R2A+PEG; n=10) and wet ecotypic *A. gerardii* rhizosphere samples (R2A; n=10 and R2A+PEG; n=10).

### DNA extraction, shotgun sequencing, and analyses

We extracted the microbial DNA with the E.Z.N.A. Soil DNA Kit (Omega Bio-tek, Inc., Norcross, GA, USA) following the manufacturer’s protocol. Shotgun metagenomes were sequenced from the extracted samples on the Illumina NovaSeq 6000 (Illumina, San Diego, CA, United States), with a 150 bp paired-end sequencing strategy, with Nextera DNA Flex for library preparation and S1 flow cell. We used the program ‘iu-filer-quality-minoche’ [38] to process the short metagenomic reads, and the quality-filtered short reads were assembled into longer contiguous sequences (contigs) using MetaHit [39] with a minimum contig length of 1000 bp. We then identified open reading frames (ORFs) in the contigs, and recruited metagenomic short reads to the contigs. We used CONCOCT [40] to bin the metagenomes, and used anvi’o ver 7.0 [41] to manually curate the bins into metagenome-assembled genomes (MAGs) that satisfied the conditions of >70% completion and <10% redundancy based on single copy genes. NCBI’s Cluster of Orthologous Groups (COGs) [42] was used to assign functions to the ORFs. The MAGs were assigned to taxa using the single-copy core genes of bacteria and archaea. We further compared the resolved MAGs using Average Nucleotide Identity (ANI) [43] to identify non-redundant MAGs based on 95% ANI [44].

### Phylogenetic, pathway, and pangenomic analyses

Among the resolved MAGs, there was a MAG of interest for this study: MAG-*Pseudomonas*. The selected non-redundant MAG was analyzed by Similar Genome Finder service that uses the MinHash on the Pathosystems Resource Integration Center (PATRIC) web portal [45,46]. Similar genomes deposited in public databases were obtained and used to estimate the genome distances to the MAG-*Pseudomonas*. We constructed a phylogenetic tree for the selected non-redundant MAG and 40 closely related genomes. The workflow used the PATRIC Codon Tree Service which used the amino acid and nucleotide sequences from a well-defined database of global protein families [47]. Then, we used the RAxML program [48] to construct a tree based on the pairwise differences between the aligned protein families of the selected sequences.

We used the comparative pathway tool of PATRIC to predict the metabolic pathways in our selected MAG. To compare the pathways, we selected *Pseudomonas* genomes from rhizospheres of cotton and soybean. KEGG maps and heat maps of the nitrogen metabolism pathway were generated in the PATRIC portal. We downloaded 6 closely-related *Pseudomonas* genomes from NCBI RefSeq [49] and performed pangenomic analyses using anvi’o workflow [41,50]. The workflow uses BLASTP [51] to compute amino acid level similarities between all possible ORF pairs. We then used Markov Cluster Algorithm (MCL) [52] to group ORFs into homologous gene clusters and aligned amino acid sequences in each gene cluster using MUSCLE for visualization [53]. We determined the core gene clusters of the MAG-*Pseudomonas* and the 6 additional, available *Pseudomonas* genomes, as well as the accessory gene cluster of MAG-*Pseudomonas*.

### Availability of data

The raw data used in this study are publicly available at NCBI under the project accession PRJNA844897. Analyzed data in the form of databases and fasta files can be found at figshare https://doi.org/10.6084/m9.figshare.20005550.

## Figures and Tables legend

Supplementary Table S1: Non-redundant MAGs and taxonomic identity.

Supplementary Table S2: Gene functions of MAG-*Pseudomonas*.

Supplementary Table S3: Pathways identified in MAG-*Pseudomonas*.

Supplementary Table S4: Shared and unique gene clusters identified in pangenomic analysis of MAG-*Pseudomonas* with 6 closely-related *Pseudomonas* genomes.

## Acknowledgement

The work is supported by the National Science Foundation under Award No. OIA-1656006 and matching support from the State of Kansas through the Kansas Board of Regents. This study was also funded by the United States Department of Agriculture, National Institute of Food and Agriculture (USDA NIFA), under the Award Number: 2020-67019-31803. We appreciate the Kansas Medical Center Genomics Core for the metagenomic data generated. We are thankful to the National Institutes of Health (NIH) grant supports - Kansas Intellectual and Developmental Disabilities Research Center (NIH U54 HD 090216), the Molecular Regulation of Cell Development and Differentiation – COBRE (P30 GM122731-03) - the NIH S10, High-End Instrumentation Grant (NIH S10OD021743) and the Frontiers CTSA grant (UL1TR002366), at the University of Kansas Medical Center, Kansas City, KS 66160.

## Author’s contributions

S.S. conceptualized, conducted the experiments, performed the data analysis, and wrote the manuscript. A.K., K.W., and E.H. conducted the experiments. Q.R. and B.F. analyzed the data. M.G., A.J., and L.J. reviewed the writing for the manuscript. S.L. conceptualized, performed the data analysis, supervised, was responsible for resource acquisitions, and wrote the manuscript. All the authors contributed to the article and approved the submitted version.

## Ethics approval and consent to participate

Not Applicable.

## References

1. Borrelli P, Robinson DA, Panagos P, Lugato E, Yang JE, Alewell C, et al. Land use and climate change impacts on global soil erosion by water (2015-2070). Proc Natl Acad Sci U S A. 2020;117:21994–2001.

2. Epa US, OAR. Climate change indicators: Drought. 2016 [cited 2022 Jan 21]; Available from: https://www.epa.gov/climate-indicators/climate-change-indicators-drought

3. Sintayehu DW. Impact of climate change on biodiversity and associated key ecosystem services in Africa: a systematic review. Ecosystem Health and Sustainability. Taylor & Francis; 2018;4:225–39.

4. Turner TR, James EK, Poole PS. The plant microbiome [Internet]. Genome Biology. 2013. Available from: http://dx.doi.org/10.1186/gb-2013-14-6-209

5. Vukanti RV. Structure and Function of Rhizobiome. Plant Microbe Symbiosis. Springer, Cham; 2020. p. 241–61.

6. Bhaduri D, Pal S, Purakayastha TJ, Chakraborty K, Yadav RS, Akhtar MS. Soil Quality and Plant-Microbe Interactions in the Rhizosphere. In: Lichtfouse E, editor. Sustainable Agriculture Reviews: Volume 17. Cham: Springer International Publishing; 2015. p. 307–35.

7. Olanrewaju OS, Ayangbenro AS, Glick BR, Babalola OO. Plant health: feedback effect of root exudates-rhizobiome interactions. Appl Microbiol Biotechnol. 2019;103:1155–66.

8. Bais HP, Weir TL, Perry LG, Gilroy S, Vivanco JM. The role of root exudates in rhizosphere interactions with plants and other organisms. Annu Rev Plant Biol. 2006;57:233–66.

9. Barea J-M, Pozo MJ, Azcón R, Azcón-Aguilar C. Microbial co-operation in the rhizosphere. J Exp Bot. 2005;56:1761–78.

10. Adl S. Rhizosphere, food security, and climate change: A critical role for plant-soil research. Rhizosphere. Elsevier BV; 2016;1:1–3.

11. Ahkami AH, Allen White R, Handakumbura PP, Jansson C. Rhizosphere engineering: Enhancing sustainable plant ecosystem productivity. Rhizosphere. 2017;3:233–43.

12. Shahzad R, Khan AL, Bilal S, Waqas M, Kang S-M, Lee I-J. Inoculation of abscisic acid-producing endophytic bacteria enhances salinity stress tolerance in Oryza sativa. Environ Exp Bot. 2017;136:68–77.

13. Compant S, Van Der Heijden MGA, Sessitsch A. Climate change effects on beneficial plant–microorganism interactions. FEMS Microbiol Ecol. Oxford Academic; 2010;73:197–214.

14. Classen AT, Sundqvist MK, Henning JA, Newman GS, Moore JAM, Cregger MA, et al. Direct and indirect effects of climate change on soil microbial and soil microbial-plant interactions: What lies ahead? [Internet]. Ecosphere. 2015. p. art130. Available from: http://dx.doi.org/10.1890/es15-00217.1

15. Alshaal T, El-Ramady H, Al-Saeedi AH, Shalaby T, Elsakhawy T, Omara AE-D, et al. The Rhizosphere and Plant Nutrition Under Climate Change. In: Naeem M, Ansari AA, Gill SS, editors. Essential Plant Nutrients: Uptake, Use Efficiency, and Management. Cham: Springer International Publishing; 2017. p. 275–308.

16. Pugnaire FI, Morillo JA, Peñuelas J, Reich PB, Bardgett RD, Gaxiola A, et al. Climate change effects on plant-soil feedbacks and consequences for biodiversity and functioning of terrestrial ecosystems. Sci Adv. 2019;5:eaaz1834.

17. Naylor D, Coleman-Derr D. Drought Stress and Root-Associated Bacterial Communities. Front Plant Sci. 2017;8:2223.

18. Monohon SJ, Manter DK, Vivanco JM. Conditioned soils reveal plant-selected microbial communities that impact plant drought response. Sci Rep. 2021;11:21153.

19. Poudel M, Mendes R, Costa LAS, Bueno CG, Meng Y, Folimonova SY, et al. The Role of Plant-Associated Bacteria, Fungi, and Viruses in Drought Stress Mitigation. Front Microbiol. 2021;12:743512.

20. Wang S, Ouyang L, Ju X, Zhang L, Zhang Q, Li Y. Survey of Plant Drought-Resistance Promoting Bacteria from Populus euphratica Tree Living in Arid Area. Indian J Microbiol. 2014;54:419–26.

21. Rolli E, Marasco R, Vigani G, Ettoumi B, Mapelli F, Deangelis ML, et al. Improved plant resistance to drought is promoted by the root-associated microbiome as a water stress-dependent trait [Internet]. Environmental Microbiology. 2015. p. 316–31. Available from: http://dx.doi.org/10.1111/1462-2920.12439

22. Lu X, Taylor AE, Myrold DD, Neufeld JD. Expanding perspectives of soil nitrification to include ammonia-oxidizing archaea and comammox bacteria. Soil Sci Soc Am J. Wiley; 2020;84:287–302.

23. Zhang J, Liu Y-X, Zhang N, Hu B, Jin T, Xu H, et al. NRT1.1B is associated with root microbiota composition and nitrogen use in field-grown rice. Nat Biotechnol. 2019;37:676–84.

24. Marín O, González B, Poupin MJ. From Microbial Dynamics to Functionality in the Rhizosphere: A Systematic Review of the Opportunities With Synthetic Microbial Communities. Front Plant Sci. 2021;12:650609.

25. Bai Y, Müller DB, Srinivas G, Garrido-Oter R, Potthoff E, Rott M, et al. Functional overlap of the Arabidopsis leaf and root microbiota. Nature. 2015;528:364–9.

26. Castrillo G, Teixeira PJP, Paredes SH, Law TF, de Lorenzo L, Feltcher ME, et al. Root microbiota drive direct integration of phosphate stress and immunity [Internet]. Nature. 2017. p. 513–8. Available from: http://dx.doi.org/10.1038/nature21417

27. Jansson JK, Hofmockel KS. The soil microbiome-from metagenomics to metaphenomics. Curr Opin Microbiol. 2018;43:162–8.

28. Gray MM, St. Amand P, Bello NM, Galliart MB, Knapp M, Garrett KA, et al. Ecotypes of an ecologically dominant prairie grass (Andropogon gerardii) exhibit genetic divergence across the U.S. Midwest grasslands’ environmental gradient [Internet]. Molecular Ecology. 2014. p. 6011–28. Available from: http://dx.doi.org/10.1111/mec.12993

29. Galliart M, Bello N, Knapp M, Poland J, St Amand P, Baer S, et al. Local adaptation, genetic divergence, and experimental selection in a foundation grass across the US Great Plains’ climate gradient [Internet]. Global Change Biology. 2019. p. 850–68. Available from: http://dx.doi.org/10.1111/gcb.14534

30. Sarkar S, Kamke A, Ward K, Rudick AK, Baer SG, Ran Q, et al. Bacterial but Not Fungal Rhizosphere Community Composition Differ among Perennial Grass Ecotypes under Abiotic Environmental Stress. Microbiol Spectr. 2022;e0239121.

31. de Raad M, Li Y, Andeer P, Kosina SM, Saichek NR. A defined medium based on R2A for cultivation and exometabolite profiling of soil bacteria. bioRxiv [Internet]. biorxiv.org; 2021; Available from: https://www.biorxiv.org/content/10.1101/2021.05.23.445362v1.abstract

32. Michel BE. Evaluation of the water potentials of solutions of polyethylene glycol 8000 both in the absence and presence of other solutes. Plant Physiol. 1983;72:66–70.

33. Meher, Shivakrishna P, Ashok Reddy K, Manohar Rao D. Effect of PEG-6000 imposed drought stress on RNA content, relative water content (RWC), and chlorophyll content in peanut leaves and roots. Saudi J Biol Sci. 2018;25:285–9.

34. Marulanda A, Barea J-M, Azcón R. Stimulation of Plant Growth and Drought Tolerance by Native Microorganisms (AM Fungi and Bacteria) from Dry Environments: Mechanisms Related to Bacterial Effectiveness [Internet]. Journal of Plant Growth Regulation. 2009. p. 115–24. Available from: http://dx.doi.org/10.1007/s00344-009-9079-6

35. Nelson SK, Oliver MJ. A Soil-Plate Based Pipeline for Assessing Cereal Root Growth in Response to Polyethylene Glycol (PEG)-Induced Water Deficit Stress. Front Plant Sci. 2017;8:1272.

36. Maritz JM, Sullivan SA, Prill RJ, Aksoy E, Scheid P, Carlton JM. Filthy lucre: A metagenomic pilot study of microbes found on circulating currency in New York City. PLoS One. 2017;12:e0175527.

37. Sarkar S, Ward K, Kamke A, Ran Q, Feehan B, Richie T, et al. Perspective: Simple State Communities to Study Microbial Interactions: Examples and Future Directions. Front Microbiol. 2022;13:801864.

38. Eren AM, Vineis JH, Morrison HG, Sogin ML. A filtering method to generate high quality short reads using illumina paired-end technology. PLoS One. 2013;8:e66643.

39. Li D, Luo R, Liu C-M, Leung C-M, Ting H-F, Sadakane K, et al. MEGAHIT v1. 0: A fast and scalable metagenome assembler driven by advanced methodologies and community practices. Methods. 2016;102:3–11.

40. Alneberg J, Bjarnason BS, de Bruijn I, Schirmer M, Quick J, Ijaz UZ, et al. Binning metagenomic contigs by coverage and composition [Internet]. Nature Methods. 2014. p. 1144–6. Available from: http://dx.doi.org/10.1038/nmeth.3103

41. Eren AM, Murat Eren A, Esen ÖC, Quince C, Vineis JH, Morrison HG, et al. Anvi’o: an advanced analysis and visualization platform for ‘omics data [Internet]. PeerJ. 2015. p. e1319. Available from: http://dx.doi.org/10.7717/peerj.1319

42. Marchler-Bauer A, Derbyshire MK, Gonzales NR, Lu S, Chitsaz F, Geer LY, et al. CDD: NCBI’s conserved domain database. Nucleic Acids Res. 2015;43:D222–6.

43. Yoon S-H, Ha S-M, Lim J, Kwon S, Chun J. A large-scale evaluation of algorithms to calculate average nucleotide identity. Antonie Van Leeuwenhoek. 2017;110:1281–6.

44. Richter M, Rosselló-Móra R. Shifting the genomic gold standard for the prokaryotic species definition. Proc Natl Acad Sci U S A. 2009;106:19126–31.

45. Ondov BD, Treangen TJ, Melsted P, Mallonee AB, Bergman NH, Koren S, et al. Mash: fast genome and metagenome distance estimation using MinHash [Internet]. Genome Biology. 2016. Available from: http://dx.doi.org/10.1186/s13059-016-0997-x

46. Davis JJ, Wattam AR, Aziz RK, Brettin T, Butler R, Butler RM, et al. The PATRIC Bioinformatics Resource Center: expanding data and analysis capabilities. Nucleic Acids Res. Oxford University Press; 2020;48:D606–12.

47. Davis JJ, Gerdes S, Olsen GJ, Olson R, Pusch GD, Shukla M, et al. PATtyFams: Protein Families for the Microbial Genomes in the PATRIC Database. Front Microbiol. 2016;7:118.

48. Stamatakis A. RAxML version 8: a tool for phylogenetic analysis and post-analysis of large phylogenies. Bioinformatics. 2014;30:1312–3.

49. O’Leary NA, Wright MW, Brister JR, Ciufo S, Haddad D, McVeigh R, et al. Reference sequence (RefSeq) database at NCBI: current status, taxonomic expansion, and functional annotation. Nucleic Acids Res. 2016;44:D733–45.

50. Eren AM, Murat Eren A, Kiefl E, Shaiber A, Veseli I, Miller SE, et al. Community-led, integrated, reproducible multi-omics with anvi’o [Internet]. Nature Microbiology. 2021. p. 3–6. Available from: http://dx.doi.org/10.1038/s41564-020-00834-3

51. Altschul SF, Gish W, Miller W, Myers EW, Lipman DJ. Basic local alignment search tool [Internet]. Journal of Molecular Biology. 1990. p. 403–10. Available from: http://dx.doi.org/10.1016/s0022-2836(05)80360-2

52. van Dongen S. Graph clustering by flow simulation. Graph Stimul. by flow Clust. PhD thesis, University of Utrecht. doi: 10.1016/j.cosrev.2007.05.001; 2000.

53. Edgar RC. MUSCLE: multiple sequence alignment with high accuracy and high throughput [Internet]. Nucleic Acids Research. 2004. p. 1792–7. Available from: http://dx.doi.org/10.1093/nar/gkh340

54. Niu X, Song L, Xiao Y, Ge W. Drought-Tolerant Plant Growth-Promoting Rhizobacteria Associated with Foxtail Millet in a Semi-arid Agroecosystem and Their Potential in Alleviating Drought Stress. Front Microbiol. 2017;8:2580.

55. Sah S, Singh R. Phylogenetical coherence of Pseudomonas in unexplored soils of Himalayan region. 3 Biotech. 2016;6:170.

56. Chandra D, Srivastava R, Glick BR, Sharma AK. Drought-Tolerant Pseudomonas spp. Improve the Growth Performance of Finger Millet (Eleusine coracana (L.) Gaertn.) Under Non-Stressed and Drought-Stressed Conditions. Pedosphere. 2018;28:227–40.

57. Achouak W, Sutra L, Heulin T, Meyer JM, Fromin N, Degraeve S, et al. Pseudomonas brassicacearum sp. nov. and Pseudomonas thivervalensis sp. nov., two root-associated bacteria isolated from Brassica napus and Arabidopsis thaliana. Int J Syst Evol Microbiol. 2000;50 Pt 1:9–18.

58. Siegele Deborah A. Universal Stress Proteins in Escherichia coli. J Bacteriol. American Society for Microbiology; 2005;187:6253–4.

59. Ghosh D, Gupta A, Mohapatra S. A comparative analysis of exopolysaccharide and phytohormone secretions by four drought-tolerant rhizobacterial strains and their impact on osmotic-stress mitigation in Arabidopsis thaliana. World J Microbiol Biotechnol. 2019;35:90.

60. Ma Y, Dias MC, Freitas H. Drought and Salinity Stress Responses and Microbe-Induced Tolerance in Plants. Front Plant Sci. 2020;11:591911.

61. Schnider-Keel U, Lejbølle KB, Baehler E, Haas D, Keel C. The sigma factor AlgU (AlgT) controls exopolysaccharide production and tolerance towards desiccation and osmotic stress in the biocontrol agent Pseudomonas fluorescens CHA0. Appl Environ Microbiol. 2001;67:5683–93.

62. Chang W-S, van de Mortel M, Nielsen L, de Guzman GN, Li X, Halverson LJ. Alginate Production by Pseudomonas putida Creates a Hydrated Microenvironment and Contributes to Biofilm Architecture and Stress Tolerance under Water-Limiting Conditions [Internet]. Journal of Bacteriology. 2007. p. 8290–9. Available from: http://dx.doi.org/10.1128/jb.00727-07

63. Ali SZ, Sandhya V, Venkateswar Rao L. Isolation and characterization of drought-tolerant ACC deaminase and exopolysaccharide-producing fluorescent Pseudomonas sp. Ann Microbiol. 2014;64:493–502.

64. Pravisya P, Jayaram KM, Yusuf A. Biotic priming with Pseudomonas fluorescens induce drought stress tolerance in Abelmoschus esculentus (L.) Moench (Okra). Physiol Mol Biol Plants. 2019;25:101–12.

65. Gulez G, Altintas A, Fazli M, Dechesne A, Workman CT, Tolker-Nielsen T, et al. Colony morphology and transcriptome profiling of Pseudomonas putida KT2440 and its mutants deficient in alginate or all EPS synthesis under controlled matric potentials. Microbiologyopen. 2014;3:457–69.

66. Woodcock SD, Syson K, Little RH, Ward D, Sifouna D, Brown JKM, et al. Trehalose and α-glucan mediate distinct abiotic stress responses in Pseudomonas aeruginosa. PLoS Genet. 2021;17:e1009524.

67. Shimada T, Katayama Y, Kawakita S, Ogasawara H, Nakano M, Yamamoto K, et al. A novel regulator RcdA of the csgD gene encoding the master regulator of biofilm formation in Escherichia coli. Microbiologyopen. 2012;1:381–94.

68. Cruz de Carvalho MH. Drought stress and reactive oxygen species: Production, scavenging and signaling. Plant Signal Behav. 2008;3:156–65.

69. Ezraty B, Gennaris A, Barras F, Collet J-F. Oxidative stress, protein damage and repair in bacteria. Nat Rev Microbiol. 2017;15:385–96.

70. Singh DP, Singh V, Gupta VK, Shukla R, Prabha R, Sarma BK, et al. Microbial inoculation in rice regulates antioxidative reactions and defense related genes to mitigate drought stress. Sci Rep. 2020;10:4818.

71. Abdelaal K, AlKahtani M, Attia K, Hafez Y, Király L, Künstler A. The Role of Plant Growth-Promoting Bacteria in Alleviating the Adverse Effects of Drought on Plants. Biology [Internet]. 2021;10. Available from: http://dx.doi.org/10.3390/biology10060520

72. Yan Y, Yang J, Dou Y, Chen M, Ping S, Peng J, et al. Nitrogen fixation island and rhizosphere competence traits in the genome of root-associated Pseudomonas stutzeri A1501. Proc Natl Acad Sci U S A. 2008;105:7564–9.

73. Hervás AB, Canosa I, Little R, Dixon R, Santero E. NtrC-dependent regulatory network for nitrogen assimilation in Pseudomonas putida. J Bacteriol. American Society for Microbiology; 2009;191:6123–35.

74. García-González V, Jiménez-Fernández A, Hervás AB, Canosa I, Santero E, Govantes F. Distinct roles for NtrC and GlnK in nitrogen regulation of the Pseudomonas sp. strain ADP cyanuric acid utilization operon. FEMS Microbiol Lett. 2009;300:222–9.

75. Setten L, Soto G, Mozzicafreddo M, Fox AR, Lisi C, Cuccioloni M, et al. Correction: Engineering Pseudomonas protegens Pf-5 for Nitrogen Fixation and its Application to Improve Plant Growth under Nitrogen-Deficient Conditions [Internet]. PLoS ONE. 2013. Available from: http://dx.doi.org/10.1371/annotation/279fe0d7-d9b1-4d05-a45a-5ff00b4606b7

76. Fox AR, Soto G, Valverde C, Russo D, Lagares A Jr, Zorreguieta Á, et al. Major cereal crops benefit from biological nitrogen fixation when inoculated with the nitrogen-fixing bacterium Pseudomonas protegens Pf-5 X940. Environ Microbiol. Wiley; 2016;18:3522–34.

77. Pankievicz VCS, Irving TB, Maia LGS, Ané J-M. Are we there yet? The long walk towards the development of efficient symbiotic associations between nitrogen-fixing bacteria and non-leguminous crops. BMC Biol. 2019;17:99.

78. Mahmud K, Makaju S, Ibrahim R, Missaoui A. Current Progress in Nitrogen Fixing Plants and Microbiome Research. Plants [Internet]. 2020;9. Available from: http://dx.doi.org/10.3390/plants9010097

79. Lyons EM, Thiel T. Characterization of nifB, nifS, and nifU genes in the cyanobacterium Anabaena variabilis: NifB is required for the vanadium-dependent nitrogenase. J Bacteriol. 1995;177:1570–5.

80. Dai Z, Guo X, Yin H, Liang Y, Cong J, Liu X. Identification of nitrogen-fixing genes and gene clusters from metagenomic library of acid mine drainage. PLoS One. 2014;9:e87976.

81. Calatrava-Morales N, Nogales J, Ameztoy K, van Steenbergen B, Soto MJ. The NtrY/NtrX System of Sinorhizobium meliloti GR4 Regulates Motility, EPS I Production, and Nitrogen Metabolism but Is Dispensable for Symbiotic Nitrogen Fixation. Mol Plant Microbe Interact. 2017;30:566–77.

82. Kramer G, Weiss V. Functional dissection of the transmitter module of the histidine kinase NtrB in Escherichia coli. Proc Natl Acad Sci U S A. 1999;96:604–9.

83. Huergo LF, Chandra G, Merrick M. P(II) signal transduction proteins: nitrogen regulation and beyond. FEMS Microbiol Rev. 2013;37:251–83.

84. Kamp A, Høgslund S, Risgaard-Petersen N, Stief P. Nitrate Storage and Dissimilatory Nitrate Reduction by Eukaryotic Microbes. Front Microbiol. 2015;6:1492.

85. Cabello P, Roldán Castillo F, Moreno-Vivián C. Nitrogen Cycle. In: Schaechter M, editor. Encyclopedia of Microbiology (Third Edition). Oxford: Academic Press; 2009. p. 299–321.

86. Wang X, Tamiev D, Alagurajan J, DiSpirito AA, Phillips GJ, Hargrove MS. The role of the NADH-dependent nitrite reductase, Nir, from Escherichia coli in fermentative ammonification. Arch Microbiol. 2019;201:519–30.

87. Robertson GP, Groffman PM. Nitrogen Transformations [Internet]. Soil Microbiology, Ecology and Biochemistry. 2015. p. 421–46. Available from: http://dx.doi.org/10.1016/b978-0-12-415955-6.00014-1

88. Meharg A. Marschner’s Mineral Nutrition of Higher Plants. 3rd edition. Edited by P. Marschner. Amsterdam, Netherlands: Elsevier/Academic Press (2011), pp. 684, US$124.95. ISBN 978-0-12-384905-2 [Internet]. Experimental Agriculture. 2012. p. 305–305. Available from: http://dx.doi.org/10.1017/s001447971100130x

89. Temple SJ, Vance CP, Stephen Gantt J. Glutamate synthase and nitrogen assimilation. Trends Plant Sci. 1998;3:51–6.

90. Miflin BJ, Habash DZ. The role of glutamine synthetase and glutamate dehydrogenase in nitrogen assimilation and possibilities for improvement in the nitrogen utilization of crops. J Exp Bot. 2002;53:979–87.

91. Mertens J, Aliyu H, Cowan DA. LEA Proteins and the Evolution of the WHy Domain. Appl Environ Microbiol [Internet]. 2018;84. Available from: http://dx.doi.org/10.1128/AEM.00539-18

92. Ciccarelli FD, Bork P. The WHy domain mediates the response to desiccation in plants and bacteria. Bioinformatics. 2005;21:1304–7.

93. Anderson D, Ferreras E, Trindade M, Cowan D. A novel bacterial Water Hypersensitivity-like protein shows in vivo protection against cold and freeze damage. FEMS Microbiol Lett. 2015;362:fnv110.

94. Kvint K, Nachin L, Diez A, Nyström T. The bacterial universal stress protein: function and regulation. Curr Opin Microbiol. 2003;6:140–5.

95. Liu P, Myo T, Ma W, Lan D, Qi T, Guo J, et al. TaTypA, a Ribosome-Binding GTPase Protein, Positively Regulates Wheat Resistance to the Stripe Rust Fungus. Front Plant Sci. 2016;7:873.

96. Weber H, Polen T, Heuveling J, Wendisch VF, Hengge R. Genome-wide analysis of the general stress response network in Escherichia coli: sigmaS-dependent genes, promoters, and sigma factor selectivity. J Bacteriol. 2005;187:1591–603.

97. Schmalisch M, Langbein I, Stülke J. The general stress protein Ctc of Bacillus subtilis is a ribosomal protein. J Mol Microbiol Biotechnol. 2002;4:495–501.

98. Hida A, Oku S, Miura M, Matsuda H, Tajima T, Kato J. Characterization of methyl-accepting chemotaxis proteins (MCPs) for amino acids in plant-growth-promoting rhizobacterium Pseudomonas protegens CHA0 and enhancement of amino acid chemotaxis by MCP genes overexpression. Biosci Biotechnol Biochem. 2020;84:1948–57.

99. Kehry MR, Dahlquist FW. The methyl-accepting chemotaxis proteins of Escherichia coli. Identification of the multiple methylation sites on methyl-accepting chemotaxis protein I. J Biol Chem. 1982;257:10378–86.

100. Muff TJ, Ordal GW. The CheC phosphatase regulates chemotactic adaptation through CheD. J Biol Chem. 2007;282:34120–8.

101. Glekas GD, Plutz MJ, Walukiewicz HE, Allen GM, Rao CV, Ordal GW. Elucidation of the multiple roles of CheD in Bacillus subtilis chemotaxis. Mol Microbiol. 2012;86:743–56.

102. Yuan W, Glekas GD, Allen GM, Walukiewicz HE, Rao CV, Ordal GW. The importance of the interaction of CheD with CheC and the chemoreceptors compared to its enzymatic activity during chemotaxis in Bacillus subtilis. PLoS One. 2012;7:e50689.

103. Yang G, Waterfield NR. The role of TcdB and TccC subunits in secretion of the Photorhabdus Tcd toxin complex. PLoS Pathog. 2013;9:e1003644.

104. Fuqua WC, Winans SC, Greenberg EP. Quorum sensing in bacteria: the LuxR-LuxI family of cell density-responsive transcriptional regulators. J Bacteriol. 1994;176:269–75.

105. Li Z, Nair SK. Quorum sensing: how bacteria can coordinate activity and synchronize their response to external signals? Protein Sci. 2012;21:1403–17.

106. Rascio N, La Rocca N. Biological Nitrogen Fixation. In: Jørgensen SE, Fath BD, editors. Encyclopedia of Ecology. Oxford: Academic Press; 2008. p. 412–9.

107. Takai K. The Nitrogen Cycle: A Large, Fast, and Mystifying Cycle. Microbes Environ. 2019;34:223–5.

108. Zhuang L, Li Y, Wang Z, Yu Y, Zhang N, Yang C, et al. Synthetic community with six Pseudomonas strains screened from garlic rhizosphere microbiome promotes plant growth. Microb Biotechnol. 2021;14:488–502.

109. Sitaraman R. Pseudomonas spp. as models for plant-microbe interactions. Front Plant Sci. 2015;6:787.

110. Castellano-Hinojosa A, Strauss SL. Insights into the taxonomic and functional characterization of agricultural crop core rhizobiomes and their potential microbial drivers. Sci Rep. 2021;11:10068.

