## Supplementary material for "Culturomics of *Andropogon gerardii* rhizobiome revealed nitrogen transforming capabilities of stress-tolerant *Pseudomonas* under drought conditions": Table 1

|  | MAG-*Pseudomonas* |
| --- | --- |
| GC-content | 61.11% |
| N-50 | 146,692 bp |
| Number of contigs | 99 |
| Percent Completion | 100% |
| Percent redundancy | 1.41% |
| Total length | 6,777,975 bp |
